## Supporting Information for "Systematic Design of Pulse Dosing to Eradicate Persister Bacteria"

*SI-1 APPENDIX A. Derivation of eqn. (7)*

Consider pulse dosing with antibiotic switched on ($C \neq 0$) and off ($C = 0$) at times $t_0, t_1, t_2, t_3, \ldots$ and with respective pulse durations $t_{\text{on}}, t_{\text{off}}$. Then, eqn. (3) implies

$$\mathbf{x}(t_1) = \exp(\mathbf{A}_{\text{on}} t_{\text{on}})\, \mathbf{x}(t_0)$$

$$\mathbf{x}(t_2) = \exp(\mathbf{A}_{\text{off}} t_{\text{off}})\, \mathbf{x}(t_1) = \exp(\mathbf{A}_{\text{off}} t_{\text{off}}) \exp(\mathbf{A}_{\text{on}} t_{\text{on}})\, \mathbf{x}(t_0)$$

$$\mathbf{x}(t_3) = \cdots = \exp(\mathbf{A}_{\text{on}} t_{\text{on}}) \exp(\mathbf{A}_{\text{off}} t_{\text{off}}) \exp(\mathbf{A}_{\text{on}} t_{\text{on}})\, \mathbf{x}(t_0)$$

$$\vdots$$

$$\mathbf{x}(t_{2\ell})$$

$$= \underbrace{\left(\overbrace{\exp(\mathbf{A}_{\text{off}} t_{\text{off}}) \exp(\mathbf{A}_{\text{on}} t_{\text{on}})}^{\mathbf{M}}\right) \ldots \left(\overbrace{\exp(\mathbf{A}_{\text{off}} t_{\text{off}}) \exp(\mathbf{A}_{\text{on}} t_{\text{on}})}^{\mathbf{M}}\right)}_{\ell} \mathbf{x}(t_0)$$

$$= \mathbf{M}^{\ell} \mathbf{x}(t_0), \quad \ell = 0, 1, 2, \ldots$$

and

$$\mathbf{x}(t_{2\ell+1})$$

$$= \exp(\mathbf{A}_{\text{on}} t_{\text{on}}) \underbrace{\left(\overbrace{\exp(\mathbf{A}_{\text{off}} t_{\text{off}}) \exp(\mathbf{A}_{\text{on}} t_{\text{on}})}^{\mathbf{M}}\right) \ldots \left(\overbrace{\exp(\mathbf{A}_{\text{off}} t_{\text{off}}) \exp(\mathbf{A}_{\text{on}} t_{\text{on}})}^{\mathbf{M}}\right)}_{\ell} \mathbf{x}(t_0)$$

$$= \exp(\mathbf{A}_{\text{on}} t_{\text{on}})\, \mathbf{M}^{\ell}(t_0), \quad \ell = 0, 1, 2, \ldots$$

Therefore

$$c(t_{2\ell}) = \mathbf{c}^{\top} \mathbf{M}^{\ell} \mathbf{x}(t_0) \tag{SI-1}$$

$$c(t_{2\ell+1}) = \mathbf{c}^{\top} \exp(\mathbf{A}_{\text{on}} t_{\text{on}})\, \mathbf{M}^{\ell} \mathbf{x}(t_0) \tag{SI-2}$$

where $\mathbf{c}^{\top} = [1 \quad 1]$.

The above two equations suggest that peaks and dips of $c(t)$ at times $t_{2\ell}$ and $t_{2\ell+1}$, respectively, increase or decrease at the same rate, governed by the eigenvalues $\lambda_1, \lambda_2$ of the matrix

$$\mathbf{M} \overset{\text{def}}{=} \exp(\mathbf{A}_{\text{off}} t_{\text{off}}) \exp(\mathbf{A}_{\text{on}} t_{\text{on}}) \tag{SI-3}$$

as

$$c(t_{2\ell}) = \mathbf{c}^{\top} \mathbf{M}^{\ell} \mathbf{x}(t_0) = \mathbf{c}^{\top} \mathbf{P} \mathbf{\Lambda}^{\ell} \mathbf{P}^{-1} \mathbf{x}(t_0) \tag{SI-4}$$

and similarly for $c(t_{2\ell+1})$.

Now, because $a \approx 0$, the eigenvalues, $\rho_1, \rho_2$ of

$$\mathbf{A} \overset{\text{def}}{=} \begin{bmatrix} K_n & b \\ a & K_p \end{bmatrix} \tag{SI-5}$$

in eqn. (3) are approximately

$$\boxed{\rho_1 \approx K_n, \;\; \rho_2 \approx K_p} \tag{SI-6}$$

When the antibiotic is on, then $b \approx 0$, and

$$K_{n,\text{on}} < K_{p,\text{on}} < 0 \tag{SI-7}$$

with $K_{p,\text{on}}$ closer to 0 than $K_{n,\text{on}}$, because $K_{p,\text{on}}$ refers to persisters, in contrast to $K_{n,\text{on}}$, which refers to normal cells that get killed much faster (e.g. see Table 1). In addition, the corresponding eigenvectors of $\mathbf{A}_{\text{on}}$ are $\mathbf{w}_{1,\text{on}} = [1 \quad 0]^{\top}$, $\mathbf{w}_{2,\text{on}} = [0 \quad 1]^{\top}$. Therefore,

$$\exp(\mathbf{A}_{\text{on}} t_{\text{on}}) = \begin{bmatrix} \exp(K_{n,\text{on}} t_{\text{on}}) & 0 \\ 0 & \exp(K_{p,\text{on}} t_{\text{on}}) \end{bmatrix} \tag{SI-8}$$

Similarly, when the antibiotic is off, it follows that

$$K_{p,\text{off}} < 0 < K_{n,\text{off}} \tag{SI-9}$$

because the normal cell subpopulation grows, whereas persister cells decline due to their returning to the state of normal cells (e.g. see Table 1). In addition, the corresponding eigenvectors of $\mathbf{A}_{\text{off}}$ are $\mathbf{w}_{1,\text{off}} = [1 \quad 0]^{\top}$, $\mathbf{w}_{2,\text{off}} = [1 \quad \xi]^{\top}$ where $\xi = (K_p - K_n)/b$. Therefore

$$\exp(\mathbf{A}_{\text{off}} t_{\text{off}}) = \exp(K_{n,\text{off}} t_{\text{off}}) \, \mathbf{w}_{1,\text{off}} \mathbf{z}_{1,\text{off}}^{\top} + \exp(K_{p,\text{off}} t_{\text{off}}) \, \mathbf{w}_{2,\text{off}} \mathbf{z}_{2,\text{off}}^{\top} \tag{SI-10}$$

where $\mathbf{z}_{1,\text{off}}^{\top} = [1 \quad -1/\xi]$, $\mathbf{z}_{2,\text{off}}^{\top} = [1 \quad 1/\xi]$ are the rows of the inverse modal matrix.

Combination of the last two equations (SI-10) and (SI-8) with the above definition of $\mathbf{M}$ in eqn. (SI-3) yields

$$\mathbf{M} = \begin{bmatrix} \exp(K_{n,\text{off}} t_{\text{off}} + K_{n,\text{on}} t_{\text{on}}) & X \\ 0 & \exp(K_{p,\text{off}} t_{\text{off}} + K_{p,\text{on}} t_{\text{on}}) \end{bmatrix} \tag{SI-11}$$

where the actual form of $X$ in the above eqn. (SI-11) does not affect the eigenvalues of **M,** which are

$$\boxed{\lambda_1 = \exp(K_{n,\text{off}} t_{\text{off}} + K_{n,\text{on}} t_{\text{on}}), \;\; \lambda_2 = \exp(K_{p,\text{off}} t_{\text{off}} + K_{p,\text{on}} t_{\text{on}})} \tag{SI-12}$$

For $\mathbf{M}^{\ell}$ to decline as $\ell$ increases, it is necessary and sufficient that both $0 < \lambda_1 < 1$ and $0 < \lambda_2 < 1$, which is equivalent to

$$K_{n,\text{off}} t_{\text{off}} + K_{n,\text{on}} t_{\text{on}} < 0 \Leftrightarrow \boxed{\frac{t_{\text{off}}}{t_{\text{on}}} < -\frac{K_{n,\text{on}}}{K_{n,\text{off}}}} \tag{SI-13}$$

and

$$K_{p,\text{off}} t_{\text{off}} + K_{p,\text{on}} t_{\text{on}} < 0 \Leftrightarrow \frac{t_{\text{off}}}{t_{\text{on}}} > -\frac{K_{p,\text{on}}}{K_{p,\text{off}}} \tag{SI-14}$$

in view of the inequalities in eqns. (SI-7) and (SI-9).

The inequality in eqn. (SI-14) is trivially satisfied, given eqns. (SI-7) and (SI-9). Therefore, it is the inequality in eqn. (SI-13) that characterizes the feasible region for $t_{\mathrm{off}}/t_{\mathrm{on}}$.

---

### *SI-2 APPENDIX B. Optimal rate of decline for bacterial population peaks characterized by eqn. (9)*

Eqn.  implies that successive peaks of $c(t)$ at times $t_{2\ell}$ (Fig 1) are characterized as

$$c(t_{2\ell}) = p_1 \lambda_1^{\ell} + p_2 \lambda_2^{\ell},\ \ell = 0, 1, 2, \ldots \tag{SI-15}$$

where $\lambda_1, \lambda_2$ are the eigenvalues of $\mathbf{M}$ and $p_1, p_2$ are coefficients depending on the model parameters and initial conditions. Each of the two terms $\lambda_1^{\ell},\ \lambda_2^{\ell}$ in eqn. (SI-15) corresponds to a decline ratio from time point $t_{2\ell}$ to $t_{2(\ell+1)}$. For values of $t_{\text{off}}/t_{\text{on}}$ satisfying eqn. (SI-13), which guarantees $0 < \lambda_1 < 1,\ 0 < \lambda_2 < 1$ hence decline of successive peaks, the larger of $\lambda_1, \lambda_2$ captures the slower of the two modes of decline ratio from peak to peak. The two planes characterized by the inequalities in eqns. (SI-13) and (SI-14) written as equalities (cf. Fig 2) intersect at the straight line $\lambda_1 = \lambda_2$ which yields

$$\boxed{\frac{t_{\text{off}}}{t_{\text{on}}} = \frac{K_{p,\text{on}} - K_{n,\text{on}}}{K_{n,\text{off}} - K_{p,\text{off}}} > 0} \tag{SI-16}$$

as exemplified in  for the parameter values in Table 1.

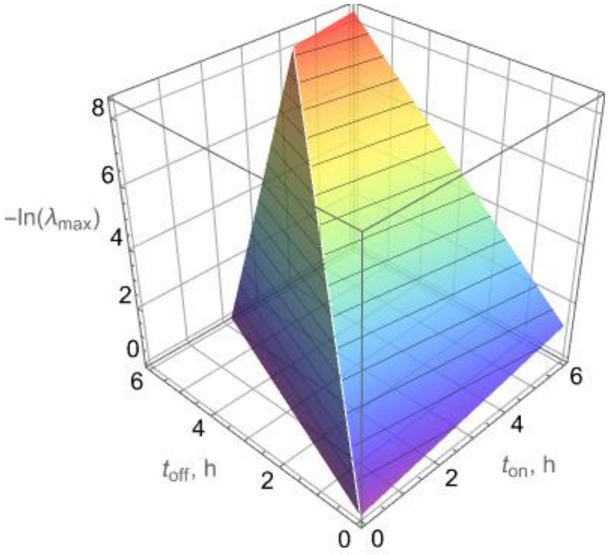


SI Fig 1. Larger (slower) of the two eigenvalues $\lambda_1, \lambda_2$ of the matrix $\mathbf{M}$ as a function of $t_{\text{on}}, t_{\text{off}}$ for the values of $K_{n,\text{on}}$, $K_{p,\text{on}}$, $K_{n,\text{off}}$, $K_{p,\text{off}}$ in Table 1. The crease line where the two planes intersect corresponds to eqn.  and characterizes the highest peak-to-peak decline ratio at discrete time points $t_{2\ell} \stackrel{\text{def}}{=} \ell(t_{\text{on}}, +t_{\text{off}})$.

The peak-to-peak decline ratio for each mode at discrete time points $t_{2\ell} \stackrel{\text{def}}{=} \ell(t_{\text{on}}, +t_{\text{off}})$ can be easily converted to actual time rate:

$$\lambda^{\ell} = \exp(\ell \ln(\lambda)) = \exp\left(\frac{t_{2\ell}}{t_{\text{on}} + t_{\text{off}}} \ln(\lambda)\right) \tag{SI-17}$$

which, combined with eqn.  immediately implies actual-time decline rates

$$k_1 = \frac{\ln(\lambda_1)}{t_{\text{on}} + t_{\text{off}}} = \frac{K_{n,\text{off}} t_{\text{off}} + K_{n,\text{on}} t_{\text{on}}}{t_{\text{on}} + t_{\text{off}}},\ k_2 = \frac{\ln(\lambda_2)}{t_{\text{on}} + t_{\text{off}}} = \frac{K_{p,\text{off}} t_{\text{off}} + K_{p,\text{on}} t_{\text{on}}}{t_{\text{on}} + t_{\text{off}}} \tag{SI-18}$$

or corresponding time constant

$$\tau_1 = \frac{1}{k_1},\ \tau_2 = \frac{1}{k_2} \tag{SI-19}$$

as indicated in Fig 11.

Note that the values of $k_1, k_2$ or $\tau_1, \tau_2$ depend on the ratio $\frac{t_{\text{off}}}{t_{\text{on}}}$ rather than on individual values of $t_{\text{on}}, t_{\text{off}}$.

To capture with a single term both modes of peak-to-peak decline corresponding to $\lambda_1, \lambda_2$ in eqn. (SI-15), one can use the geometric average of $k_1, k_2$ from eqn. (SI-18), i.e.

$$k \stackrel{\text{def}}{=} \sqrt{k_1 k_2} = \frac{\sqrt{\ln(\lambda_1)\ln(\lambda_2)}}{t_{\text{on}}+t_{\text{off}}} = \frac{\sqrt{(K_{n,\text{off}}t_{\text{off}}+K_{n,\text{on}}t_{\text{on}})(K_{p,\text{off}}t_{\text{off}}+K_{p,\text{on}}t_{\text{on}})}}{t_{\text{on}}+t_{\text{off}}} \tag{SI-20}$$

or, equivalently the geometric average of $\tau_1, \tau_2$ from eqn. (SI-19), both of which yield eqn. (9).

To characterize the maximum of the overall decline rate in eqn. (SI-20), first observe that

$$k = \frac{\sqrt{(K_{n,\text{off}}x+K_{n,\text{on}})(K_{p,\text{off}}x+K_{p,\text{on}})}}{1+x} \tag{SI-21}$$

where $x \stackrel{\text{def}}{=} \frac{t_{\text{off}}}{t_{\text{on}}}$. Then the setting $dk/dx = 0$ immediately yields

$$\left(\frac{t_{\text{off}}}{t_{\text{on}}}\right)_{\text{opt}} = \frac{2K_{n,\text{on}}K_{p,\text{on}}-K_{n,\text{on}}K_{p,\text{off}}-K_{n,\text{off}}K_{p,\text{on}}}{2K_{n,\text{off}}K_{p,\text{off}}-K_{n,\text{on}}K_{p,\text{off}}-K_{n,\text{off}}K_{p,\text{on}}} \tag{SI-22}$$

which is eqn. (11).

---

### *SI-3 APPENDIX C. Estimation of $K_{n,off}$, $K_{n,on}$ from data in Fig 5*

Assuming that persister cells are initially a tiny minority in the time-growth experiment (antibiotic off), the standard cell-balance equation is

$$\frac{dc}{dt} = K_{n,\mathrm{off}} c(t) \left(1 - \frac{c(t)}{c_{\mathrm{max}}}\right) \Leftrightarrow c(t) = c_0 \exp\left(K_{n,\mathrm{off}} t\right) \frac{1}{1 + \frac{c_0}{c_{\mathrm{max}}}\left(\exp\left(K_{n,\mathrm{off}} t\right) - 1\right)} \qquad \text{(SI-23)}$$

Application to the data shown in Fig 5(a) yields

| | Estimate | Standard Error |
|---|---|---|
| $K_{n,\mathrm{off}}$ | 1.35 | 0.09 |
| $c_0$ | $2.4 \times 10^7$ | $0.3 \times 10^7$ |
| $c_{\mathrm{max}}$ | $1.4 \times 10^9$ | $0.1 \times 10^9$ |

In the time-kill experiment, eqn. (SI-23) written for a declining population before persister cells take over becomes

$$\frac{dc}{dt} = K_{n,\mathrm{on}} c(t) \Leftrightarrow c(t) = c_0 \exp\left(K_{n,\mathrm{on}} t\right) \qquad \text{(SI-24)}$$

Application to the data shown in Fig 5(b) yields

| | Estimate | Standard Error |
|---|---|---|
| $K_{n,\mathrm{on}}$ | $-3.3$ | 1.0 |
| $\log_{10} c_0$ | 7.5 | 0.8 |

---

*SI-4 APPENDIX D. Analytical solution of eqns. (1) and (2)*

Given the eigenvalues

$$\rho_{1,2} = \frac{K_n + K_p}{2} \pm \frac{\sqrt{4ab + \left(K_n - K_p\right)^2}}{2}$$

of the matrix **A**, yields

$$c(t) = e^{\rho_1 t}\left(\underbrace{\frac{n_0 + p_0}{2} + \frac{\left(K_n - K_p\right)(n_0 - p_0)/2 + an_0 + bp_0}{\sqrt{4ab + \left(K_n - K_p\right)^2}}}_{q_1}\right)$$

$$+e^{\rho_2 t}\left(\underbrace{\frac{n_0 + p_0}{2} - \frac{\left(K_n - K_p\right)(n_0 - p_0)/2 + an_0 + bp_0}{\sqrt{4ab + \left(K_n - K_p\right)^2}}}_{q_2}\right)$$

---
